# Defensive and offensive behaviors in social interaction settings in a Kleefstra syndrome mouse model

**DOI:** 10.1101/2022.02.03.478985

**Authors:** Alejandra Alonso, Anumita Samanta, Jacqueline van der Meij, Liz van den Brand, Moritz Negwer, Irene Navarro Lobato, Lisa Genzel

**Affiliations:** Department of Neuroinformatics, Faculty of Science, Donders Institute for Brain, Cognition and Behaviour, Radboud University, P.O. Box 9010, 6500 GL Nijmegen, The Netherlands; Donders Institute for Brain, Cognition and Behaviour, RadboudUMC

**Keywords:** Kleefstra, EHMT1, Autism Spectrum Disorder, social behavior, defense, offense, aggression

## Abstract

Kleefstra syndrome in humans is characterized by general delay in development, intellectual disability and autistic features. The mouse model of this disease (*Ehmt1^+/-^*) expresses anxiety, autistic-like traits, and aberrant social interactions with non-cagemates. To investigate how *Ehmt1*^+/-^ mice behave with unfamiliar conspecifics, we allowed adult, male animals to freely interact for 10 minutes in a neutral, novel environment within a host-visitor setting. In 17 out of 74 trials there were defensive and offensive behaviors. Our key finding was that *Ehmt1*^+/-^ mice displayed defensive postures, attacking and biting; in contrast, wild-type (WT) interacting with other WT did not enact such behaviors. Further, if there was a fight between an *Ehmt1*^+/-^ and a WT mouse, it was always the *Ehmt1*^+/-^ who initiated these behaviors.

## Introduction

Kleefstra syndrome is a neurodevelopmental disorder, which is caused by haploinsufficiency of the euchromatic histone methyltransferase 1 (*EHMT1*) gene (Kleefstra et al., 2006). This gene is expressed highly in the developing neural system, acting as an epigenetic factor (Shinkai & Tachibana, 2011). In humans, this syndrome is characterized by developmental delays, distinct facial features, intellectual disability, and autistic traits (Kleefstra et al., 2006; Vermeulen et al., 2017). Around 95,7% of Kleefstra patients are diagnosed with autism spectrum disorder (Vermeulen et al., 2017), characterized by repetitive behaviors, poor verbal social communication, and deficits in social skills. Antisocial behaviors such as anger and emotional outbursts have also been reported in these patients (Kleefstra et al., 2006; Kleefstra et al., 2005; Kleefstra et al., 2009; Stewart & Kleefstra, 2007), with parents reporting occasional emotional outbursts in 46.7% of patients (Haseley, Wallis, & DeBrosse, 2021).

Drosophila models for Kleefstra syndrome target the *EHMT* ortholog (Kramer et al., 2011), producing specific loss of function mutants. These flies developed abnormal dendrite branching, expressing a decreased number of dendrite ends. At a locomotor level, larvae had altered crawling paths, without affecting total path length. Habituation, the attenuation of a response over time to a repeated stimulus, was also altered in *EHMT* mutant flies. This deficit was expressed as decreased habituation to the light-off jump reflex, which is the extension of legs and flight that is initiated upon sudden darkness. In wild-type (WT) flies this response was almost completely abolished within ten light-off events, while in *EHMT* mutants the reflex was reduced but still present in around 50% of the light-off events. In regards to social behavior, WT males were able to suppress courtship immediately after being exposed to a non-receptive female. This suppression was significantly decreased in *EHMT* mutants at short- and long-term tests, indicating a deficit in social memory (Kramer et al., 2011).

The Kleefstra mouse model, *Ehmt1*^+/-^ mice, reproduce most of the syndrome’s core symptoms (Balemans et al., 2010). *Ehmt1*^+/-^ mice have a shorter skull and nose, with their eyes positioned wide apart (Balemans et al., 2014). They have been described to display increased anxiety and reduced exploratory drive (Balemans et al., 2010). Deficits in fear extinction learning, novel object recognition and spatial object recognition have been reported in single-trial experiments (Balemans et al., 2010). However, in spatial learning studies involving multi-trial experiences and in pattern separation assays, *Ehmt1*^+/-^ mice can outperform WTs (Benevento et al., 2017; Schut et al., 2020).

At the neural network level, this mouse model suffers from hippocampal synaptic dysfunctions, such as reduced dendritic branching and spine density (Balemans et al., 2014; Balemans et al., 2010), delay in maturation of parvalbumin GABAergic interneurons in sensory cortical areas (Negwer et al., 2020), as well as irregular cortical network bursting patterns in rodent (Martens et al., 2016) and human cultures in vitro (Frega et al., 2019). Each of these could lead to disabilities in learning and in executive functions such as short-term memory and social behavior. Interestingly, an increased neurogenesis in the dentate gyrus of the hippocampus has been reported (Benevento et al., 2017).

In human Kleefstra Syndrome patients, caretakers report occasional emotional outbursts in 46.7% of the patients (Haseley et al., 2021), yet this feature has not been studied in mice. In social settings *Ehmt1*^+/-^ mice have been documented to respond in aberrant ways, from an absence of a response to inappropriate or indiscriminate approaches when confronted with unfamiliar mice (Balemans et al., 2010). In a ten-minute social play session, the time spent by two non-cagemates *Ehmt1*^+/-^ males socially engaging was on average briefer compared to WT males (Balemans et al., 2010). To measure sociability, the three-chamber task can be used, where normally WT mice first prefer but then habituate to an unfamiliar mouse placed in one of the chambers. The time spent by WT animals in the chamber with the unfamiliar mouse declines during the latter half of the session. Interestingly in *Ehmt1*^+/-^ mice this decline is not seen, which could be indicative of persistent behavior (Balemans et al., 2010).

In this study we aimed to further describe the social behavior of the *Ehmt1*^+/-^ mouse model. Social cognition is the ability of an individual to interpret social cues and react appropriately to them (Bicks, Koike, Akbarian, & Morishita, 2015). For example, the decision if and how to approach an unfamiliar individual, requires integration of the internal state of the individual – such as experience, current motivation, state of arousal – with the current state of the environment as well as any social cues from the unfamiliar individual (Chen & Hong, 2018). Such intricate orchestration of factors is dependent on the prefrontal cortex, integrating signals from hippocampal region CA2 (Stevenson & Caldwell, 2014) and CA3 (Finlay et al., 2015), hypothalamus (Anderson, 2016), and amygdala (Chen & Hong, 2018).

To this end, for the social interaction task, we adapted an existing host-visitor task (Avale et al., 2011; de Chaumont et al., 2012; Granon, Faure, & Changeux, 2003). The host mouse was placed into an open field box, to freely roam or explore for 20 min. Afterwards the visitor, an unfamiliar, non-cagemate of the same sex, was also placed within the same environment and animals were allowed to freely interact for ten minutes. The key difference to other social interaction tasks, is that the first mouse – the host – initially is alone in the novel environment before a second mouse – the visitor – is added. In our adapted version we did not socially isolate the animals before the task since it led to extreme aggressiveness (data not shown). Additionally, the period for social play was extended from four to ten minutes, since we were expecting abnormal social interactions and we wanted to provide enough time for the animals to do so.

We measured social and explorative behaviors for the duration of the encounter. *Ehmt1*^+/-^ and WT mice were combined in all possible configurations, so that each mouse was at least once a host and once a visitor, and interacted at least once with both genotypes. Each time an animal entered a trial, it was always in a novel environment.

We observed defensive and offensive behaviors in a task that normally does not elicit these among WTs. The antisocial behavior was only seen in cases when there was an *Ehmt1^+/-^ mouse* involved. Further, in such interactions between WT and Ehmt1+/-, four out of four times the *Ehmt1*^+/-^ animal initiated the offense.

## Methods

### Subjects

Ten male *Ehmt1*^+/+^ [Wild-Type (WT)] and ten male *Ehmt1^+/-^ mice* were part of this experiment. Animals were bred in-house, 12–16 weeks of age at the start of behavioral training, group housed in heterogenous genotype groups (except one cage, see below) and had ad libitum access to food and water. Mice were maintained on a 12hr/12hr light/dark cycle and tested during the light period. In compliance with Dutch law and Institutional regulations, all animal procedures were approved by the Centrale Commissie Dierproeven (CCD) and conducted in accordance with the Experiments on Animal Act. Project number 2016-014 and protocol number 2016-014-030, approved by the national CCD and local animal welfare body at Radboud University, the Netherlands.

The animals were divided into two cohorts of animals, one composed by three cages with 13 animals (8 WT, 5 *Ehmt1^+/-^*), and the second cohort by three cages with seven animals (2 WT, 5 *Ehmt1^+/-^*). All but one cage was composed of mice of both genotypes, creating heterogeneous groups from two to five animals. The difference in number of mice per cage was due to the fact that all cage mates were also littermates, and to avoid stress by remixing or isolating, the original groups from the in-house breeding facility were maintained. No persistent fighting was observed among cagemates.

### Habituation

All animals were extensively habituated to the experimenters by handling for a period of two weeks before habituation to the experimental environment. By the end of the second week the animals freely climbed upon the experimenters’ hands and showed no sign of distress when picked up by the experimenters. Pick-ups were performed by tail until the third day of handling, from there on animals were picked up by cupping (see examples at https://www.genzellab.com/#/animal-handling/).

Mice were habituated to the experimental environment, an empty square box (75 cm × 75 cm x 40 cm) made of lacquered wood, with no deliberate cues in sight. Habituation was composed of two sessions, in the first all animals from the same cage were placed together in the box for 30 minutes. The following session each animal was left to freely roam or explore on its own for ten minutes. The next time the animal would enter the box would be during the experimental phase.

### Experimental environment

Two boxes were used simultaneously in the behavioral room, each box had distinctive colored floors, and every day the cues surrounding and inside the box were changed (Fig. 1A). Cues were composed of everyday materials (paper, fabric, tape) and were attached to the walls inside of the maze in a two-dimensional manner. Walls surrounding the box also were equipped with cues, which were bulkier, to provide three-dimensional cues. Additionally, each novel environment was paired with a particular smell, provided by a cotton pad with scented oil glued within the box, out of the animal’s reach.

**Figure 1.**
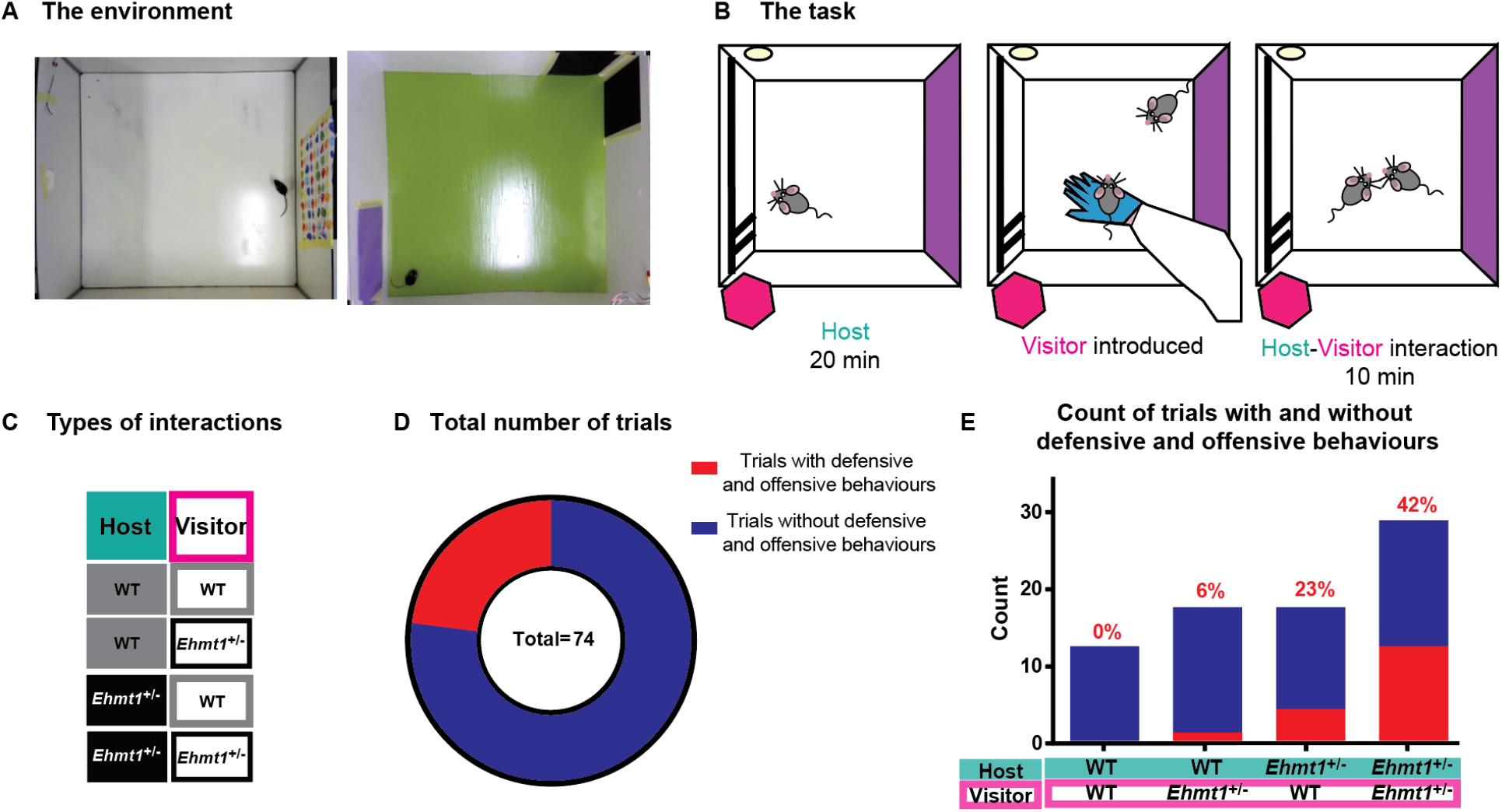
Study Design. **A.** Example of two environments. The setup consisted of a 75 × 75 cm × 40 cm square box equipped daily with new intra- and extra-maze cues, as well as a particular smell. **B.** Task. The host mouse is placed in a novel environment to roam freely for 20 min. After, an unfamiliar, non-cagemate visitor, is introduced to the same environment and left for ten minutes. Colors and stripes inside the maze represent the intra-maze cues, and the pink hexagon represents a 3D extra-maze cue. The yellow circle within the maze represents a scented cotton pad. **C.** Types of host-visitor interactions based on genotype, introducing the color code for following figures. Filled color represents the host and its genotype, grey for WT and black for *Ehmt1*^+/-^, border color represents the genotype of the visitor. Animals experienced up to one trial a day either as a host (green) or as a visitor (pink). **D.** Total number of trials and the fraction of trials that had DO-behaviors (red) or not (blue). **E.** Count of trials with DO-behaviors out of the total number of trials in each type of interaction. Red number above indicates percentage of trials with DO-behaviors within each sample. Legend under each bar represents the genotype of the host-visitor interaction. It is noticeable that defensive and offensive behaviors only occurred if an *Ehmt1*^+/-^ mouse was present (Chi-Square_3_=12.64 p=0.0055).

Video recordings were achieved with a Logitech HD Webcam C270 clamped onto the ceiling of the experimental room. Light level in the room was ~35 lux.

### The social interaction task

Before the beginning of each trial, the floor of the box was thoroughly cleaned with 70% ethanol. The trial began by placing a mouse, the host, in the experimental environment and left to roam freely. At the 20-minute mark, a non-cagemate, the visitor, was introduced to the environment, and the host-visitor pair were left to interact for ten minutes. This specific pair of animals would not be part of another trial within the same day, but could face each other later in opposite roles on other experimental days, all interactions can be seen in the interaction matrix in supplementary figure 1. Each of these animals would also act as host or visitor with different animals in successive days, until all animals had experienced both roles. Behavioral experimenters were blind to the genotype of the animal, and only intervened in case that the host-visitor pair engaged in a fight that resulted in a distress vocalization from one of the mice, in which case the trial was terminated and the animals were placed back in their homecages.

Videos were manually scored in duplicate by blinded observers, and the results of both observations were averaged.

### Statistical analysis

All behavioral data were analyzed using IBM SPSS. Data concerning exploratory behavior were analyzed using repeated-measures ANOVAs with time spent in different behaviors as the within-subject variable and genotype (*Ehmt1*^+/-^ vs. WT) and role (host or visitor) as the between-subject variable. Data concerning social interaction over sessions were analyzed using repeated-measures ANOVA with genotype (*Ehmt1*^+/-^ vs. WT) and session (1, 2, 3) as the between-subject variables. Data concerning social interaction over sessions for a single genotype were analyzed using univariate analysis with session as the between-subject variable since not all subjects completed all sessions. Data concerning interaction time in trials with defensive and offensive behaviors and those without, were analyzed using independent sample t-tests. Posterior analysis was performed using independent sample t-tests.

One outlier was removed from figure 3D, WT host - *Ehmt1*^+/-^ visitor, where time until first encounter was 124 seconds, which was more than 7 standard deviations away from the sample.

## Results

Kleefstra syndrome patients display a general delay in development, intellectual disability and autistic features (Stewart & Kleefstra, 2007). These features include stereotypical movements, impulsive behaviors, emotional outbursts, poor verbal social communication, as well as deficits in social cognition (Chevallier, Kohls, Troiani, Brodkin, & Schultz, 2012; Haseley et al., 2021; Kleefstra et al., 2005; Vermeulen et al., 2017). To investigate social behavior in the *Ehmt1*^+/-^ mouse model of the syndrome, we used a social interaction task (Fig. 1B) with unfamiliar non-cagemates, in host and visitor roles. We performed 74 trials of social interaction among *Ehmt1*^+/-^ and WT mice (Fig. 1C, D), consisting of 20 minutes of initial exploration of an open novel environment (Fig. 1A) by a host, followed by ten minutes of shared occupancy of the box (social play) with a visitor (Fig. 1B, C). We measured time spent by each of the animals exploring the environment and socially interacting. Surprisingly, 17 out of 74 host-visitor interactions showed some level of defensive and offensive behaviors (DO-behaviors) (Fig. 1D), from tail rattling to biting attacks, and 7 out of the 17 trials with such behaviors had to be terminated early due to persistent fighting. Upon analysis it became clear that these behaviors exclusively occurred if there was an *Ehmt1*^+/-^ mouse in the pair (Fig. 1E) (Chi-Square_3_=12.64 p=0.0055).

Analysis was performed for every trial, as well as for each individual animal (Supplementary Figures 2 and 3). Overall, the results remained the same in both types of analysis.

### Spatial Exploration

First, we aimed at characterizing the general activities of each animal in the box. The social interaction task started with the host animal in the open field box for 20 minutes, where the host had a chance to explore the box and the cues placed inside. The cues and the smell of the box were changed daily to increase novelty and stimulate exploration (Fig. 1A). The time spent sitting in the corners of the box or exploring the cues and walls (i.e., spatial exploration) was scored and compared between genotype (Fig. 2A). No differences were observed among *Ehmt1*^+/-^ and WT hosts during these initial 20 minutes. Hosts of both genotypes spent more time exploring the environment than sitting in the corners as seen in the main effect of behavior (ANOVA, genotype F_1,72_=1.4; p=0.237, behavior F_1,72_=6.9; p=0.011, behavior*genotype F_1,72_=0.7; p=0.396).

**Figure 2.**
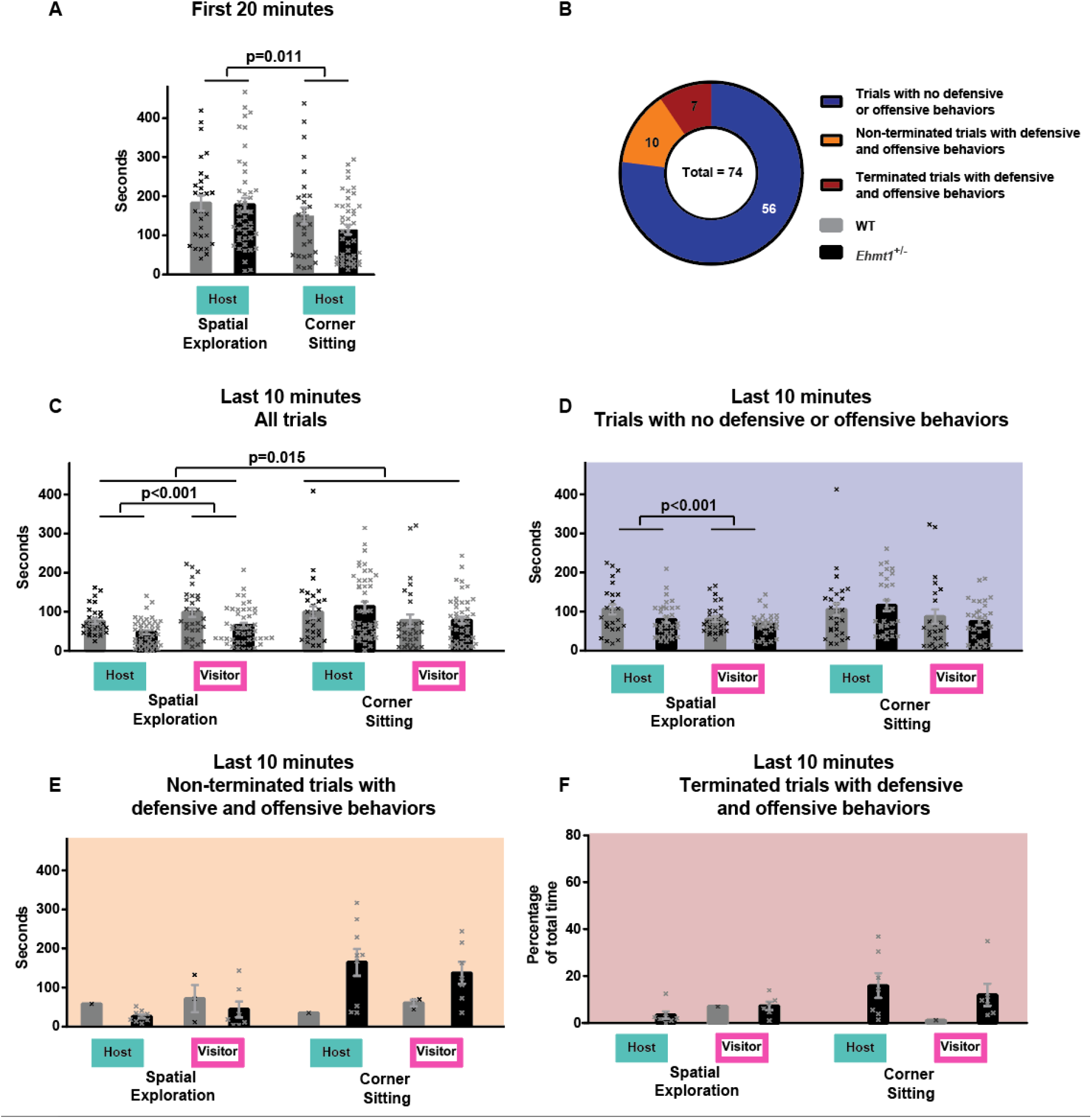
Average time spent corner sitting and spatial exploration the open-field for. **A**, host during the first 20 minutes of the task (ANOVA, genotype F_1,72_=1.4; p=0.237, behavior F_1,72_=6.9; p=0.011, behavior*genotype F_1,72_=0.7; p=0.396). **B**, Total number of trials and the fraction of trials that had DO-behaviors features which ended up in terminating a trial (dark red) or not (orange). On blue, trials that had no DO-behaviors features. **C** Host and visitor during the last ten minutes of the task, for all 74 trials (ANOVA behavior F_1,144_=6.12, p=0.015, genotype F_1,144_=3.15, p=0.21, role F_1,144_=0.84, p=0.006, behavior*genotype F_1,144_=5.4, p=0.021, behavior*role F_1,144_=8.37, p=0.004, behavior*genotype*role F_1,144_=0.07, p=0.795). **D**, host and visitor during the last ten minutes of the task, only for trials that had no DO-behaviors (behavior F_1,110_=1.502, p=0.223, genotype F_1,110_=2.304, p=0.132, role F_1,110_=0.361, p=0.361, behavior*genotype F_1,110_=1.120, p=0.292, behavior*role F_1,110_=7.108, p=0.009, gene*role F_1,110_=1.743, p=0.190, behavior*gene*role F_1,110_=0.034, p=0.853). **E**, host and visitor during the last ten minutes of the task, only for trials that had DO-behaviors but were not terminated (behavior F_1,16_=2.589, p=0.127, genotype F_1,16_=1.547, p=0.231, role F_1,16_=0.062, p=0.806, behavior*genotype F_1,16_=4.736, p=0.045, behavior*role F_1,16_=0.069, p=0.797, role*gene F_1,16_=0.171, p=0.685, behavior*gene*role F_1,16_=0.216, p=0.648). **F**, host and visitor during the last ten minutes of the task, only for trials that had DO-behaviors and had to be terminated early. Interaction time is shown as percentage of total interaction time, taking into account the length of each terminated trial. Crosses correspond to individual trials. Error bar corresponds to SEM.

During the last 10 minutes the host-visitor interaction took place, and as mentioned before, out of the 74 interactions, 17 had DO-behaviors, and out of these, 10 were 10-minute trials, while 7 had to be terminated early (Fig. 2B). Below we analyze separately these different instances.

For all 74 interactions (Fig. 2C), irrespective if they had DO-behaviors or not, there was a significant difference among behavior as well as for role (host or visitor). Additionally, there was a significant interaction among behavior and genotype, as well as among behavior and role (ANOVA behavior F_1,144_=6.12, p=0.015, genotype F_1,144_=3.15, p=0.21, role F_1,144_=0.84, p=0.006, behavior*genotype F_1,144_=5.4, p=0.021, behavior*role F_1,144_=8.37, p=0.004, behavior*genotype*role F_1,144_=0.07, p=0.795). Analyzing sitting behaviors of *Ehmt1*^+/-^ and WT mice, irrespective of their role, showed no significant differences (two-sample t_146_=0.59, p=0.962), while for spatial exploration there was a non-significant trend showing that *Ehmt1*^+/-^ animals tended to explore slightly less than WT (two-sample t_146_=3.99, p=0.062). When analyzing the host and visitor roles irrespective of genotype, there was no difference in the time spent corner sitting, (two-sample t_146_=2.52, p=0.098), while the time spent exploring was significantly greater for the visitor than the host (two-sample t_146_=2.23, p<0.001).

Next, analyzing only the trials with no DO-behaviors (Fig. 2D), there was a significant interaction for behavior and role (ANOVA behavior F_1,110_=1.502, p=0.223, genotype F_1,110_=2.304, p=0.132, role F_1,110_=0.361, p=0.361, behavior*genotype F_1,110_=1.120, p=0.292, behavior*role F_1,110_=7.108, p=0.009, gene*role F_1,110_=1.743, p=0.190, behavior*gene*role F_1,110_=0.034, p=0.853). When analyzing sitting behaviors of the host and visitor roles irrespective of genotype in trials with no DO-behaviors, there was no effect (two-sample t_112_=2.316 p=0.275), while the time spent exploring was significantly greater for the visitor than the host (two-sample t_112_=2.031, p<0.001), similar to what we had seen for all trials.

In non-terminated trials with DO-behaviors (Fig. 2E), analysis showed that there was a significant interaction among behavior and genotype (ANOVA behavior F_1,16_=2.589, p=0.127, genotype F_1,16_=1.547, p=0.231, role F_1,16_=0.062, p=0.806, behavior*genotype F_1,16_=4.736, p=0.045, behavior*role F_1,126_=0.069, p=0.797, role*gene F_1,16_=0.171, p=0.685, behavior*gene*role F_1,16_=0.216, p=0.648). Analyzing sitting behaviors of *Ehmt1*^+/-^ and WT mice, irrespective of their role, there was a significant difference, with *Ehmt1*^+/-^ mice spending more time sitting than WT mice (two-sample t_18_=2.134 p=0.018), and no differences for spatial exploration (two-sample t_18_=1.552 p=0.561).

In terminated trials (Fig. 2F), there were no instances of WT hosts, and only one instance of a WT visitor, so a full statistical model could not be run in this data set.

To test if differences were seen depending on the presence of DO-behaviors, we analyzed the behavior in trials with no DO-behaviors and in non-terminated with DO-behaviors (terminated trials were not included due to them not having a total of 600 seconds within the environment). There was a significative interaction of behavior and genotype, as well as marginal effect (p=0.055) in the interaction of DO-behaviors and genotype (ANOVA DO-behaviors F_1,126_=1.135, p=0.289, behavior F_1,126_=3.477, p=0.065, genotype F_1,126_=1.233, p=0.269, role F1,126=0.003, p=0.956, behavior* DO-behaviors F_1,126_=1.349, p=0.248, behavior*genotype F_1,126_=5.523, p=0.020, behavior*role F_1,126_=1.022, p=0.314, behavior* DO-behaviors *genotype F_1,126_=3.038, p=0.084, behavior*DO-behaviors*role F_1,126_=0.269, p=0.605, behavior*genotype*role F_1,126_=0.240, p=0.625, behavior*DO-behaviors*genotype*role F_1,126_=0.147, p=0.702, DO-behaviors*genotype F_1,126_=3.755, p=0.055, DO-behaviors*role F_1,126_=0.309, p=0.579, genotype*role F_1,126_=0.751, p=0.388, DO-behaviors*genotype*role F_1,126_=0.022, p=0.883).

In sum, during the initial 20-min exploration period, hosts spent longer exploring the environment than sitting, but there were no differences among genotypes. During the last 10 minutes, visitors spent more time exploring than the hosts, which was expected from animals entering a novel environment. In non-terminated trials with DO-behaviors, *Ehmt1*^+/-^ mice spent longer sitting than WTs. Overall, genotype did not influence exploratory and sitting behaviors in trials without DO-behaviors, while on trials with DO-behaviors *Ehmt1*^+/-^ animals spent longer sitting than WTs.

### Social exploration

Next, we focused on social behavior. Social interaction was scored as direct contact between the two animals, and again the analysis is subdivided for trials with no DO-behaviors (Fig. 3A), non-terminated trials with DO-behaviors (Fig. 3B) and terminated trials with DO-behaviors (Fig. 3C). For the trials with no DO-behaviors, time spent socially interacting was significantly longer if there was an *Ehmt1*^+/-^ animal as a host, but there was no effect of the genotype of the visitor and no significant interaction (ANOVA genotype_host F_1,53_=17.957, p<0.001, genotype_visitor F_1,53_=2.385, p=0.128, genotype_host * genotype_visitor F_1,53_=0.046 p=0.832).

**Figure 3.**
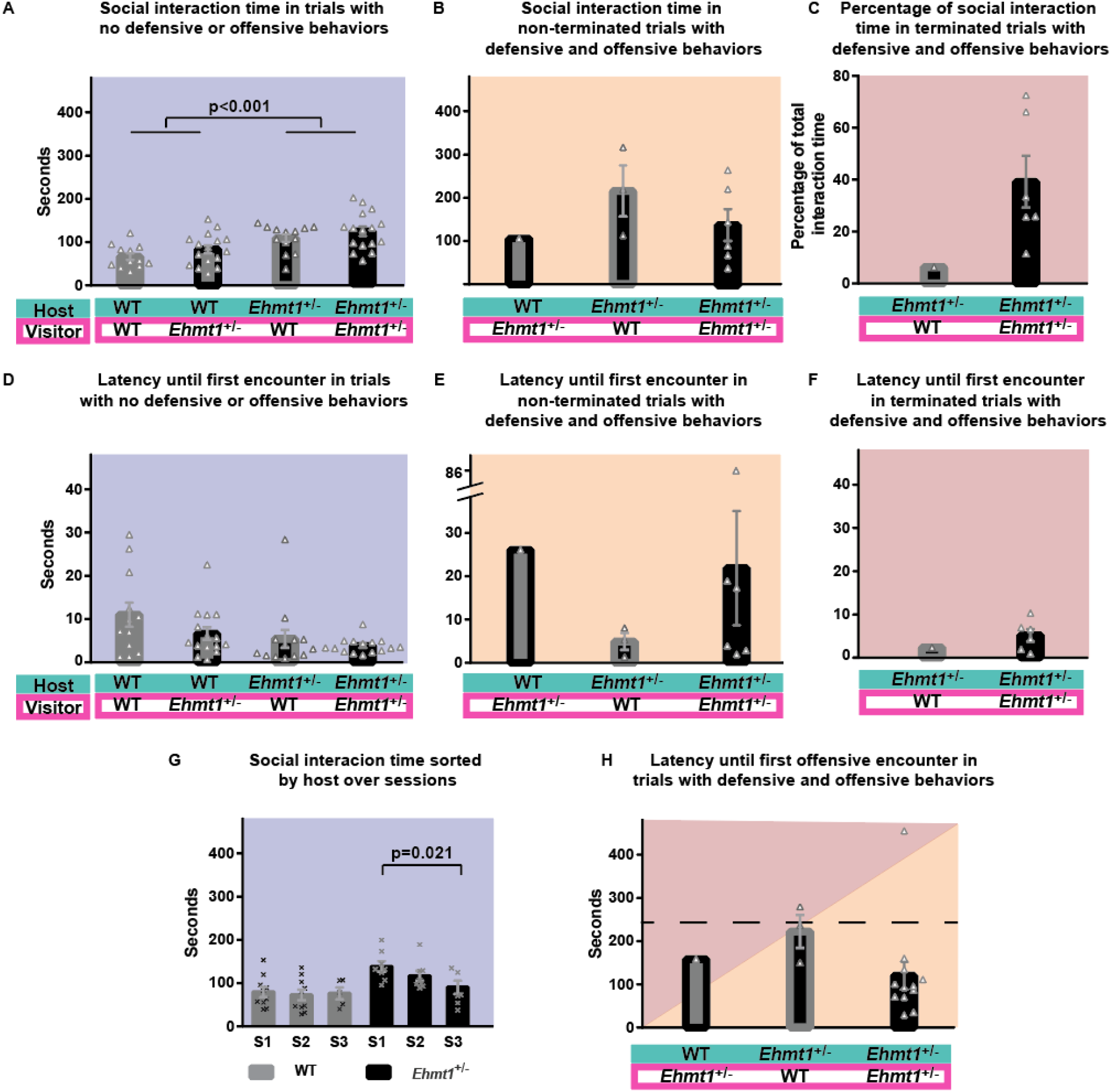
Social interaction time. **A.** Average social interaction time out of a total of ten minutes in trials with no DO-behaviors (ANOVA genotype_host F_1,53_=17.957, p<0.001, genotype_visitor F_1,53_=2.385, p=0.128, genotype_host * genotype_visitor F_1,53_=0.046 p=0.832). **B.** Average social interaction time out of a total of ten minutes in trials with DO-behaviors that were not terminated (ANOVA, genotype_visitor F_1,7_=0.749, p=0.415). **C.** Percentage of social interaction time out of the total interaction in trials with DO-behaviors that were terminated early. **D.** Average latency until first encounter, from the moment that the visitor is introduced to the maze for trials with no DO-behaviors (ANOVA genotype_host F_1,53_=3.127, p=0.083, genotype_visitor F_1,53_=0.018, p=0.895, genotype_host * genotype_visitor F_1,53_=0.282, p=0.598). **E.** Average latency until first encounter, from the moment that the visitor is introduced to the maze for non-terminated trials with DO-behaviors (ANOVA F_1,7_=0.749, p=0.415). **F.** Average latency until first encounter, from the moment that the visitor is introduced to the maze in trials with DO-behaviors that were terminated early. **G.** Average social interaction time of the host during the first 3 sessions (S1, S2 and S3) for trials with no DO-behaviors (ANOVA genotype_host F_1,42_=11.752, p=0.001, session F_2,42_=2.430, p=0.100, session*genotype_host F_2,42_=1.958, p=0.154) (ANOVA session F_2,20_=4.734; p = 0.021). Each animal had maximum one trial per day, as a host or as a visitor. **H.** Average latency until first aggressive encounter, for all trials with DO-behaviors (ANOVA genotype_visitor F_1,14_=0.597, p=0.452). Dashed line marks the 4-minute mark, which is the length of a trial in the original task. Crosses or triangles correspond to individual trials. Error bar corresponds to SEM. Background color indicates if the trials had no DO-behaviors (blue), had DO-behaviors but were non-terminated (orange), and had DO-behaviors and were terminated (red). Interaction times were longer if there was an Ehmt1+/- animal involved, but this decreased over consecutive trials.

In non-terminated trials with DO-behaviors (Fig. 3B), there was only one instance of a WT host with *Ehmt1*^+/-^ visitor, therefore analysis was performed only for *Ehmt1*^+/-^ hosts. The time spent interacting in non-terminated trials was not influenced by the genotype of the visitor (ANOVA, genotype_visitor F_1,7_=0.749, p=0.415).

In terminated trials with DO-behaviors (Fig. 3C), there was only one instance of *Ehmt1*^+/-^ host with a WT visitor, therefore statistical analysis of interaction time nor latency (Fig. 3F) were not possible. The average length of terminated trials was 320 seconds (an uninterrupted trial is 600 seconds), and the social interaction time was measured as a percentage out of the total length of each particular terminated trial. In *Ehmt1*^+/-^ - *Ehmt1*^+/-^ interactions, animals spent on average 40% of the trial interacting (Fig. 3C), around 20% of the time sitting in corners, while exploring was the less preferred activity (Fig 2F).

Latency until the first social encounter was analyzed for trials with no DO-behaviors (Fig. 3D), and there was no difference among genotype (ANOVA genotype_host F_1,53_=3.127, p=0.083, genotype_visitor F_1,53_=0.018, p=0.895, genotype_host * genotype_visitor F_1,53_=0.282, p=0.598) although there was a non-significant trend (p=0.083) for *Ehmt1*^+/-^ hosts approaching unfamiliar mice faster than WTs. In non-terminated trials with DO-behaviors (Fig. 3E), there was only one instance of a WT host with *Ehmt1*^+/-^ visitor, so analysis was performed only for *Ehmt1*^+/-^ hosts. Latency to approach an unfamiliar non-cagemate was not influenced by the genotype of the visitor (ANOVA F_1,7_=0.749, p=0.415).

Each animal experienced a maximum of one trial per day, either as a host or a visitor, and on following days they could perform the task again, until every animal experienced the role of host and visitor at least once. To analyze if their behavior changed over consecutive trials on different days, we evaluated the time spent by each host engaging in social interaction, but now split for consecutive trials (Fig. 3G). We could see an effect of the genotype of the host, but no effect over consecutive sessions nor interaction (session F_2,42_=2.430, p=0.100, genotype_host F_1,42_= 11.752, p=0.001, session*genotype_host F_2,42_=1.958, p=0.154). A trend was noticeable where the amount of time interacting decreased over time in the Ehmt1+/- animals. When analyzing the time spent interacting over consecutive days only for *Ehmt1*^+/-^ hosts, there was a significant decrease over time (ANOVA session F_2,20_=4.734; p = 0.021), while when performing the same analysis for WT hosts, no differences could be seen across sessions (session F_2,22_=0.09, p=0.910, Fig. 3G). For all terminated and non-terminated trials with DO-behaviors, the latency until the first DO-behavior interaction can be seen in Fig. 3H, and no statistical differences were observed depending on the visitor (ANOVA genotype_visitor F_1,14_=0.597, p=0.452). If there was a DO-behavior interaction involving an *Ehmt1*^+/-^ animal, 88% of the time it was before four minutes had elapsed, marked as a dashed line in figure 3H. The 4-minute mark is used as a reference since it indicates the length of a trial in the original task by Avale et al (Avale et al., 2011).

The average time of social interaction with an *Ehmt1*^+/-^ animal present for terminated and non-terminated trials was not different (two-sample t_15_=1.550, p=0.204), however interaction time in trials with DO-behaviors was longer than in trials with no DO-behaviors (two-sample t_60_=1.759, p<0.001).

In sum, *Ehmt1*^+/-^ animals spent significantly more time interacting with unfamiliar animals. The main differences were always observed depending on the genotype of the host, but overall, there seemed to be a cumulative effect where there was a linear change from two wild-types, to mixed interactions, to finally two *Ehmt1*^+/-^ with more time interacting.

### Defensive-offensive scale

In trials with DO-behaviors, levels of defense and offense were scored using a defensive-offensive scale per individual (Fig. 4). The scale ranged from 0 to 3, with 0 being no defense, 1 if the animal stands on their hind paws and/or rattles its tail, 2 when animals engage in fight, and 3 when the fight reaches a point where one mouse emits a distress vocalization (screech). In this case the offender is scored with 3 while the offended animal depending on its behavior, is scored from 0 to 2 (if the offended mouse fights back the score would be 2, if it just takes defensive postures the score would be 1, and if the mouse doesn’t defend itself, the score would be 0). Once an animal in a trial reached score 3, the host-visitor pair were separated by the experimenter and the trial was terminated, to avoid stress and harm to the animals. Analyzing the defensive-offensive scale values revealed that when there are two *Ehmt1*^+/-^ animals defensive-offensive levels were similar for host and visitor, but when there is an *Ehmt1*^+/-^ - WT pairing, the *Ehmt1*^+/-^ is always the offender.

**Figure 4.**
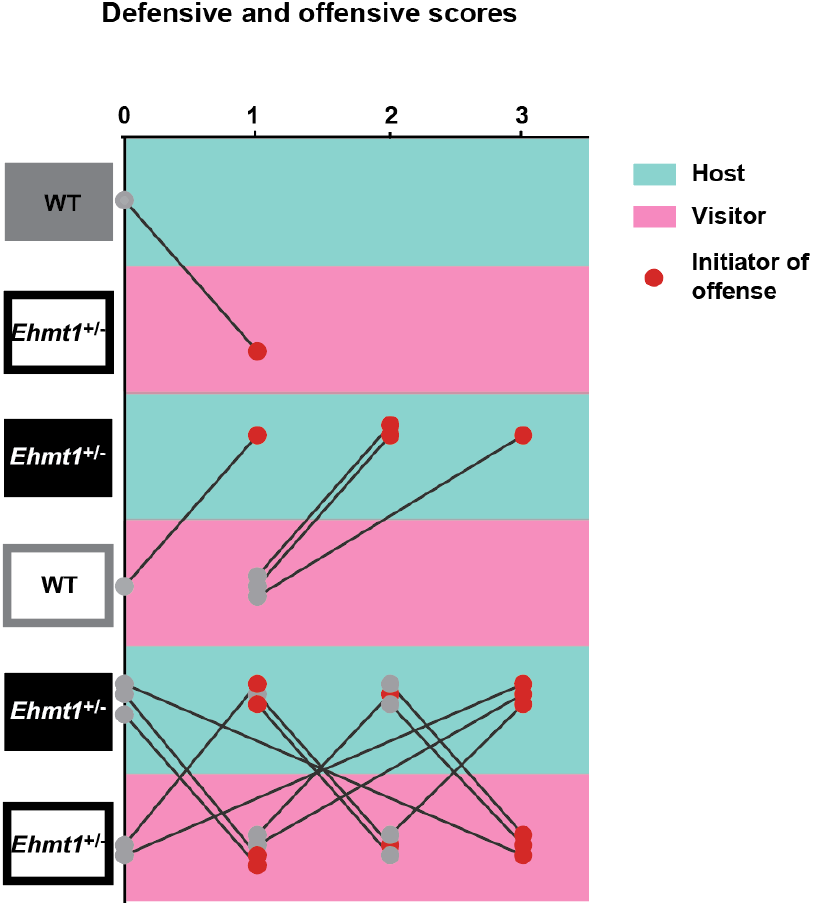
Defensive and offensive scores. Individual defensive and offensive scores for host-visitor interactions. Scale: 0: No defense, 1: Defensive posture (hind paw stand) and/or snake rattle, 2: Fight (biting), 3: Terminated trial due to one of the animals generating a distress sound. Filled circles correspond to values of individual trials, red filled circles to the animal that initiated the offense. Lines connect each host-visitor pairing. Background color indicates if it corresponds to host or visitor.

The initiator of the offense was also identified in all such interactions, in *Ehmt1*^+/-^ - *Ehmt1*^+/-^ interactions both the host and visitor were equally likely to initiate the first offensive behavior (in 6 instances the initiator was the host, and in 6 instances the initiator was the visitor). In *Ehmt1^+/-^* - WT pairings, it was always the *Ehmt1*^+/-^ animal who initiated the offense.

In sum DO-behaviors were only observed when *Ehmt1*^+/-^ animals were involved, and in the case of *Ehmt1*^+/-^ and WT pairings, it was always the *Ehmt1*^+/-^ animal that initiated the offense.

## Discussion

In this study we used a social interaction task adapted from Granon et al (2003), in a Kleefstra syndrome mouse model (*Ehmt1*^+/-^ mice). The task has not been reported to elicit antisocial behaviors among C57BL6/j mice (Avale et al., 2011; Besson, Suarez, Cormier, Changeux, & Granon, 2008; Granon et al., 2003), but, as we show here, elicits DO-behaviors behavior in *Ehmt1*^+/-^ mice. In our adapted version of the task (see below) a host would spend 20 minutes in a novel environment, after which a non-cagemate – the visitor – was introduced to the environment and allowed to freely interact with the host for ten minutes.

Our key finding was that *Ehmt1*^+/-^ mice displayed defensive postures, attacking and biting; in contrast, WT interacting with other WT did not enact such behaviors. Further, if there was a fight between an *Ehmt1*^+/-^ and a WT mouse, it was always the *Ehmt1*^+/-^ who initiated the offense. 43% of the *Ehmt1*^+/-^ – *Ehmt1*^+/-^ interactions, and 23% of the *Ehmt1*^+/-^ - WT interactions had DO-behaviors features. Every *Ehmt1*^+/-^ animal from this study was part of at least one defensive/offensive interaction (Supp. Fig. 1). In interactions with no DO-behaviors, the time spent interacting was longer if there was an *Ehmt1*^+/-^ mouse involved.

Emotional outbursts are described by caretakers in 46.7% of Kleefstra patients, a symptom which affects quality of life for both patients and caretakers (Haseley et al., 2021). Previous studies investigating the behavior of *Ehmt1*^+/-^ mice in social paradigms (Balemans et al., 2010) have not reported any behavior resembling emotional outbursts. To understand the underlying neurobiological mechanisms affected by Kleefstra syndrome, animal models are crucial. Just as important is the use of adequate behavioral tests that capture key symptoms that may create patient and caregiver suffering. The task used in this study revealed a specific phenotype, which has not been captured in other social tasks: DO-behaviors when confronted with an unfamiliar non-cagemate in a neutral environment.

### Task

The behavioral paradigm used in our study was adapted from Granon et al (2003), where they wanted to evaluate the role of prefrontal cortex β2 nicotinic receptors in novel social interaction. To enhance social interaction during the task they created a particular host-visitor setting. The host was socially isolated for four weeks and then had 30 minutes to explore a novel environment, where the interaction would take place. Then a non-isolated, unfamiliar, non-cagemate visitor of the same sex and age joined the environment for four minutes. By isolating the host prior and allowing individual exploration of the environment, when the visitor was introduced, the host would have a high drive to socially interact and a lower drive for exploring the environment. We adapted this task for *Ehmt1*^+/-^ mice. In the current study mice were not socially isolated before the task, since in previous experiments we had observed exaggerated aggressiveness from socially isolated *Ehmt1*^+/-^ mice (data not shown), and there is evidence that housing mixed genotypes improves sociability among autistic models (Yang, Perry, Weber, Katz, & Crawley, 2011). All mice from this study lived in cages from two to four animals of mixed genotype, with the exception of one cage with two animals with the same genotype.

Another adaptation to the original task was the period of the interaction, instead of only four minutes we extended until ten, while the initial solitary exploration by the host was reduced from 30 to 20 minutes. These adjustments were to provide enough time for social interactions to occur, as it had been reported that *Ehmt1*^+/-^ mice display low exploratory drive, increased sitting time and shorter social interaction time (Balemans et al., 2010). In the present study we found no difference among genotypes in sitting or exploratory behavior during the first 20 minutes, but we did find differences for social interaction during the last 10 minutes. During the ten-minute interaction period, WTs never showed DO-behaviors towards other WTs, as we expected from previous studies (Avale et al., 2011; Besson et al., 2008; Granon et al., 2003), while *Ehmt1*^+/-^ mice did show DO-behaviors towards animals of either genotype. These defensive-offensive interactions took place before four minutes had elapsed in 88% of the cases. Therefore, the extended interaction period provided in our task was likely not the reason for the occurrence of DO-behaviors in a task not previously known to lead to such responses (Avale et al., 2011; Besson et al., 2008; Granon et al., 2003).

By eliminating a confound of increased aggressiveness due to single housing, this task unveils DO-behaviors never reported before in *Ehmt1*^+/-^ mice.

### Social behavior

Previous studies investigating the social behavior of *Ehmt1*^+/-^ mice describe aberrant interactions. Balemans et al. (2010), performed a social play task as well as the three-chamber task with this mouse model. In the social play task juveniles were isolated 30 minutes prior a social play session, which took place in a clean cage for ten minutes with a WT non-cagemate of the same age and sex. They found that the time spent interacting was significantly lower in *Ehmt1*^+/-^ mice in comparison to WT mice, contradicting our results. In our study, by allowing 20 minutes of exploration prior social play, instead of isolating 30 minutes prior, hosts had enough time to explore the environment. As seen in Avale et al. (2011), upon introduction of a novel animal, the focus of exploring a novel environment would be shifted towards a novel individual, which may have driven the increased interaction time for *Ehmt1*^+/-^ mice compared to Balemans et al. (2010). Even though shorter social interaction was reported in the social play task, in the same study described above prolonged sociability was found in *Ehmt1*^+/-^ mice in the three-chamber task (Balemans et al., 2010). The three-chamber task consists of one chamber with a non-cagemate in a small, wire cage, a chamber with an empty wire cage, and a central empty chamber connecting the first two. In a ten-minute trial, WT mice decreased the time spent interacting with the contained unfamiliar animal during the last five minutes in comparison to the first five minutes. In contrast, *Ehmt1*^+/-^ mice spent roughly the same time interacting in the first and last five minutes of the trial. Similar to our findings in which *Ehmt1*^+/-^ mice spend longer time interacting, the mice in Balemans et al’s study persisted to interact across the length of the session (Balemans et al., 2010).

Social impairments are key symptoms in autism spectrum disorder (Chevallier et al., 2012), and in rodents such impairments can be expressed as a decreased social response or as the opposite, an exaggerated or inadequate response. Several mice models for autism show social impairments like low social motivation, increased self-grooming, and impaired nest building (Moy et al., 2009; for a detailed list, see supplementary materials of Varghese et al., 2017), while some strains have been reported to show increased aggression or DO-behaviors behaviors (Burrows et al., 2015; Grayton, Missler, Collier, & Fernandes, 2013). DO-behaviors has been linked to autism in mice (Burrows et al., 2015; Grayton et al., 2013), and in these models’ DO-behaviors is related to abnormal response inhibition. In some transgenic lines the aggression phenotype has been rescued by normalizing excitatory synaptic transmission (Miyamoto et al., 2017; Selimbeyoglu et al., 2017), or by treating with serotonin and dopamine antagonists (Burrows et al., 2015).

An imbalance in the ratio of excitatory and inhibitory synapses within the cortical, mnemonic and emotional circuits is thought to contribute to the occurrence of diseases with deficits in social behaviors such as autism (Antoine, Langberg, Schnepel, & Feldman, 2019; Gogolla et al., 2009; Rubenstein & Merzenich, 2003; Selten, van Bokhoven, & Nadif Kasri, 2018), which has also been reported in the *Ehmt1*^+/-^ mouse model (Negwer et al., 2020), and could be underlying the increased DO-behaviors that we report in this current study.

Animals lacking cortical β2 subunit nicotinic receptor (Avale et al., 2011) also show an imbalance in excitatory synaptic transmission. The animals in Avale’s study displayed increased time in social contact and expressed exaggerated approaches towards novel mice, behavior which could be rescued by re-expressing the β2 subunit in the prelimbic cortex (Avale et al., 2011). This behavior is similar to what we observe in our study, increased social interaction time when there is an *Ehmt1*^+/-^ mouse involved. When evaluating latency to approach an unfamiliar mouse, although non-significant (p=0.083), our results suggest that *Ehmt1*^+/-^ mice are faster in approaching than WTs, which could be interpreted as an exaggerated approach towards an unfamiliar mouse.

Another example is the *Stxbp1*^+/-^ mouse model. *Stxbp1* gene encodes a synaptic protein essential for neurotransmitter release, and patients suffering from this mutation exhibit epilepsy, intellectual disability, autism and aggressive behaviors (Miyamoto et al., 2017). *Stxbp1*^+/-^ mice were not aggressive in a ten-minute interaction task within an open field, but did show increased contact time. However, in a resident intruder test, where a WT intruder is introduced to a socially isolated resident mouse’s homecage for ten minutes, *Stxbp1*^+/-^ mice were highly aggressive towards the intruder. Our task is in some ways similar to the resident intruder paradigm, since the host spends 20 minutes on their own in the novel environment, giving the animal a chance to mark territory. Nevertheless, in *Ehmt1*^+/-^ WT pairings, no matter who was the host, it was always the *Ehmt1*^+/-^ animal who initiated DO-behaviors. *Stxbp1*^+/-^ aggressive behavior was suppressed after treating them with ampakines, which potentiates excitatory transmission (Miyamoto et al., 2017).

Work in cell cultures from Kleefstra syndrome patients, as well as auditory cortex slices of *Ehmt1*^+/-^ mice (Frega et al., 2019), has shown that there is an upregulation of NMDA receptor subunit 1 in excitatory cortical neurons, which affects the excitatory-inhibitory balance. This imbalance can be rescued by inhibiting NMDA receptors (Frega et al., 2019). Similarly, studies on cortical cultures of *Ehmt1*^+/-^ mice revealed that synaptic scaling is also disturbed, a process that regulates inhibition and excitation (Benevento et al., 2016). In sensory and auditory cortices delays in maturation of parvalbumin-positive interneurons have been described in histological analysis of *Ehmt1*^+/-^ mouse brains (Negwer et al., 2020). Additionally, studies in cell cultures with Ehmt1 knock down expressed impairments in spontaneous network activity, producing a delay in cortical firing development (Martens et al., 2016).

The CA2 region of the hippocampus in rodents is necessary for social recognition, processing the what, when and where of social information (Oliva, Fernandez-Ruiz, Leroy, & Siegelbaum, 2020; Tzakis & Holahan, 2019). Inhibition (Hitti & Siegelbaum, 2014) or lesions (Stevenson & Caldwell, 2014) of CA2 in mice impaired social recognition of a co-housed littermate when compared to a novel non-cagemate, but spatial and contextual memory, as well as olfactory systems were spared. Neural network studies have identified a circuit in which CA2 promotes aggression by activating the lateral septum, which in turn disinhibits the ventrolateral subnucleus of the ventromedial hypothalamus. This nucleus is directly implicated in regulating aggression (Anderson, 2016; Leroy et al., 2018). So far, in the hippocampus the dentate gyrus (Benevento et al., 2017), hippocampal region CA1 and CA3 (Balemans et al., 2014) region, and entorhinal cortex (Gupta-Agarwal et al., 2012) have been reported to differ from WT in *Ehmt1*^+/-^ mice, but studies focused on CA2 have not been performed yet.

In sum, in our results we observe aberrant social behavior in *Ehmt1*^+/-^ mice, expressed as prolonged sociability and exaggerated approach towards unfamiliar animals. We report DO-behaviors for the first time in this mouse line, a feature which has also been reported in autism models. An imbalance in excitatory-inhibitory synapses is likely underlying these behavioral impairments.

### Habituation deficits

Another recurrent symptom in patients with autism and in autism animal models is decreased habituation, which may be also playing a role in the DO-behaviors seen in our study.

Habituation can be defined as one of the simplest forms of learning, where an individual stops responding to a stimuli that has been repeatedly presented (Rankin et al., 2009). In humans with autism spectrum disorder, sensory habituation is reduced from an early age (Guiraud et al., 2011) and neural habituation has been described in humans in the amygdala, correlating low levels of habituation to severe social impairments (Kleinhans et al., 2009).

Studies in fly have also reported that EHMT mutants display a deficit in habituation during the light-off jump reflex. WT flies cease to perform a jump reflex after being repeatedly exposed to a light-off, they habituate to the stimulus. In contrast, *EHMT* mutants were never able to fully suppress the reflex, as it was still present in around half of the events (Kramer et al., 2011). Further, studies in several fly models for intellectual disability and autism spectrum disorder have repeatedly found this habituation deficit (Fenckova et al., 2019).

Similarly, *Ehmt1*^+/-^ mice fail to habituate to a novel unfamiliar mouse in a three-chamber task (Balemans et al., 2010). In a previous study with *Ehmt1*^+/-^ animals, we exposed animals to an object exploration task for three consecutive weeks. The first week of training *Ehmt1*^+/-^ mice were exploring less than WT animals. As weeks went by, exploration time increased in *Ehmt1*^+/-^ mice, reaching the same values as WTs (Schut et al., 2020), similar to what we report in this study. *Ehmt1*^+/-^ hosts decrease the time they spent interacting after experiencing the task for a third time, reaching similar values to WTs.

A failure or a delay to habituate may contribute to increased anxiety, and increased levels of anxiety may lead to aggressiveness. Increased social interaction time has been related to anxiety in rats (File & Hyde, 1978), however in some aggressive mice strains it seems that high levels of results in lower levels of anxiety (Nyberg, Vekovischeva, & Sandnabba, 2003). The interplay of anxiety, aggression and habituation is partly modulated by interneurons in cortical areas such as anterior cingulate cortex (Chaibi, Bennis, & Ba-M’Hamed, 2021), and as discussed in the previous sections, *Ehmt1*^+/-^ mice suffer from cortical imbalances in excitatory-inhibitory signaling. Therefore, altered habituation may be linked to excitatory-inhibitory imbalances in this mouse model.

By extensively handling *Ehmt1*^+/-^ animals in this experiment the anxiety reported previously (Balemans et al., 2010) may have decreased. With less anxious animals, we observe exploration periods similar to WT animals during the first 20 minutes of solitary exploration of the environment. At social interaction level, a significant habituation effect was seen over following sessions, decreasing the interaction time up to levels comparable to WTs. However, in some individuals this decrease in anxiety may also have led to increased aggression (Nyberg et al., 2003).

Overall, this suggests that longer periods of training in *Ehmt1*^+/-^ animals may counteract their anxiety phenotype, coupled to extensive handling and mixed grouped housing, this may result in changes in behavior such as increased exploratory drive. Therefore, it is tempting to speculate that habituation deficits and delayed habituation underlies many of the symptoms seen in this syndrome.

### Conclusions

By using an adapted social interaction task that combines a novel environment with a host-visitor interaction, we were able to show aberrant social approach and defensive/offensive behaviors in the *Ehmt1*^+/-^ mouse model. These findings have two implications. Firstly, they provide a target for trials on potential treatments of emotional outbursts. Secondly, they show that this task can be used to test for features of DO-behaviors towards unfamiliar animals in other rodent models of disease.

## Acknowledgements

We would like to thank Nael Nadif Kasri for providing the animals and valuable feedback during manuscript preparation. We would also like to thank all students who have performed the social interaction task: Yarick Winmai, Niamh Hemingway and Alysha Maurmair, as well as students scoring the videos: Manon van Bentum, Esra Canki, Annie Do Tramh Anh, and Nienke Cremers.

## Funding

This work was supported by the M-GATE project, which receives funding from the European Union’s Horizon 2020 research and innovation programme under the Marie Skłodowska-Curie grant agreement No 765549, and the Branco Weiss Fellowship to Lisa Genzel.

## Supplementary Materials

**Supplementary Figure 1.**
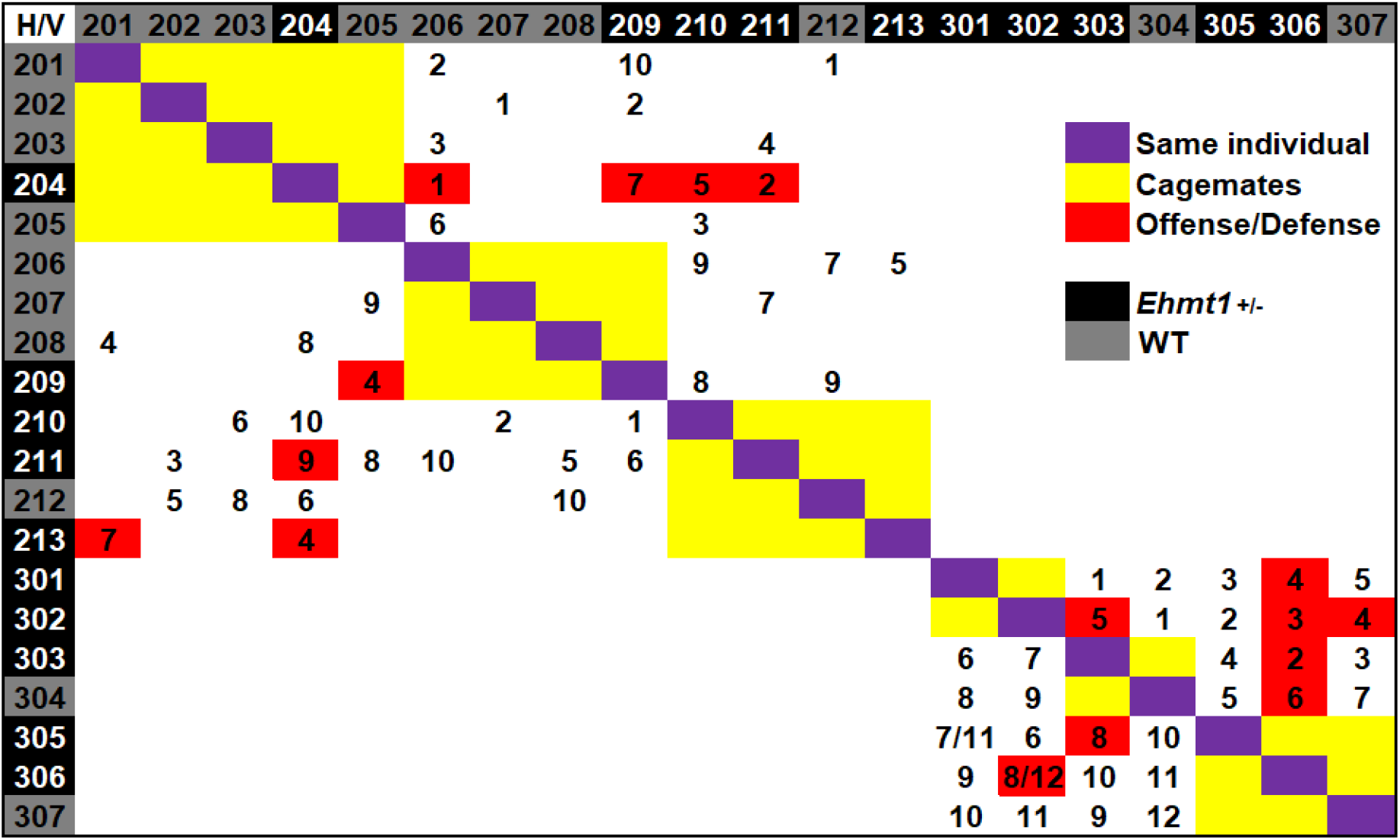
Interaction matrix. Rows correspond to host (H) and columns to visitor (V), three-digit number corresponds to animal identification. Numbers within the matrix correspond to the day on which the trial took place.

**Supplementary Figure 2.**
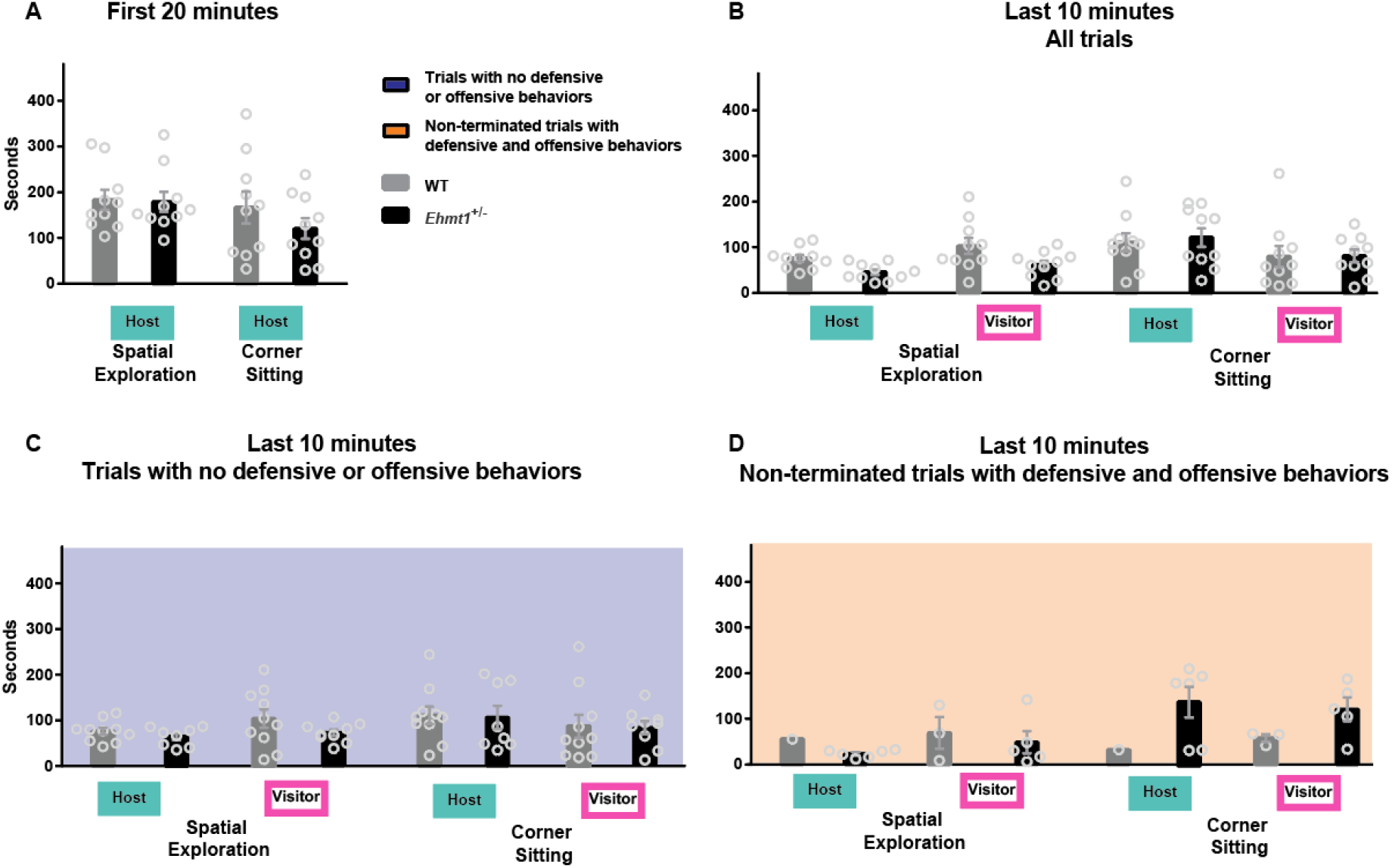
Average time spent corner sitting and spatial exploration the open-field for. **A**, host during the first 20 minutes of the task (ANOVA behaviour F_1,18_=2.23, p=0.153, genotype F_1,18_=0.89, p=0.356, behaviour*genotype F_1,18_=0.68, p=0.420). **B**, Host and visitor during the last ten minutes of the task, for all 74 trials (ANOVA behaviour F_1,36_=27.99, p<0.001, genotype F_1,36_=3.00, p=0.091, role F_1,36_=0.02, p=0.881, behaviour*genotype F_1,36_=0.14, p=0.709, behaviour*role F_1,36_=0.02, p=0.877, behaviour*genotype*role F_1,36_=2.23, p=0.143). **C**, host and visitor during the last ten minutes of the task, only for trials that had no DO-behaviors (Behavior F_1,33_=1.176, p=0.286, Genotype F_1,33_=2.853, p=0.101, Role F_1,33_=0.198, p=0.659, Behaviour*genotype F_1,33_=0.234, p=0.632, Behaviour*role F_1,33_=1.851, p=0.183, Behaviour*genotype*role F_1,33_= 0.085, p=0.773, Genotype*role F_1,33_=0.425, p=0.519). **D**, host and visitor during the last ten minutes of the task, only for trials that had DO-behaviors but were not terminated (Behavior F_1,11_=2.154, p=0.110, Genotype F_1,11_=1.364, p=0.268, Role F_1,11_=0.204, p=0.660, Behaviour*genotype F_1,11_=4.519, p=0.057, Behaviour*role F_1,11_=0.089, p=0.772, Behaviour*genotype*role F_1,11_=0.284, p=0.605, Genotype*role F_1,11_=0.118, p=0.738). Circles correspond to the average per animal. Error bar corresponds to SEM.

**Supplementary Figure 3.**
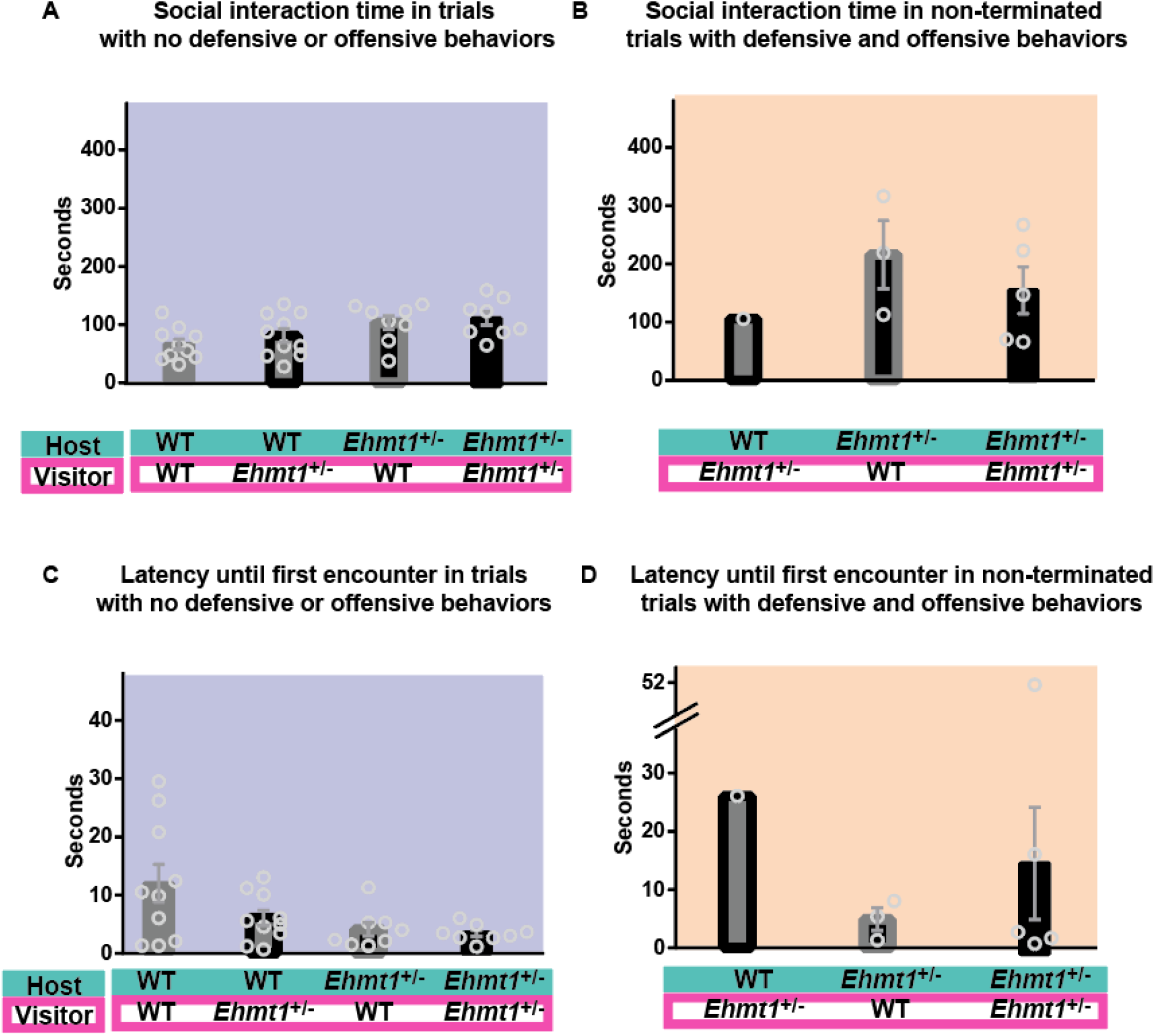
Social interaction time. **A.** Average social interaction time out of a total of ten minutes in trials with no DO-behaviors (ANOVA genotype_host F_1,32_=10.543, p=0.003, genotype_visitor F_1,32_=1.564, p=0.220, genotype_host*genotype_visitor F_1,32_=0.011, p=0.916). **B.** Average social interaction time out of a total of ten minutes in trials with DO-behaviors that were not terminated (ANOVA Genotype_host F_1,6_ =0.227, p=0.651, Gentoype_visitor F_1,6_ =0.803, p=0.405). **C.** Average latency until first encounter, from the moment that the visitor is introduced to the maze for trials with no DO-behaviors (ANOVA genotype_host F_1,32_=6.578, p=0.015, genotype_visitor F_1,32_=2.607, p=0.116, genotype_host*genotype_visitor F_1,32_=1.610, p=0.214). **D.** Average latency until first encounter, from the moment that the visitor is introduced to the maze for non-terminated trials with DO-behaviors (ANOVA Genotype_host F_1,6_ =0.294, p=0.607, Genotype_visitor F_1,6_ =0.680, p=0.441). Circles correspond to average values per individual. Error bar corresponds to SEM. Background color indicates if the trials had no DO-behaviors (blue) or had DO-behaviors but were non-terminated (orange).

## Notes

### Competing Interest Statement

The authors have declared no competing interest.

